# Selection of an Appropriate Empiric Antibiotic Regimen in Culture-Negative Hematogenous Vertebral Osteomyelitis

**DOI:** 10.1101/363549

**Authors:** Ki-Ho Park, Dong Youn Kim, Yu-Mi Lee, Mi Suk Lee, Kyung-Chung Kang, Jung Hee Lee, Seong Yeon Park, Chisook Moon, Yong Pil Chong, Sung-Han Kim, Sang-Oh Lee, Sang-Ho Choi, Yang Soo Kim, Jun Hee Woo, In-Gyu Bae, Oh-Hyun Cho

## Abstract

The aim of this study was to determine which antibiotic combinations are appropriate for culture-negative hematogenous vertebral osteomyelitis (HVO), based on the antibiotic-susceptibility pattern of organisms isolated from cases of culture-proven HVO. We conducted a retrospective chart review of adult patients with microbiologically proven HVO in five tertiary-care hospitals over a 7-year period. The appropriateness of empiric antibiotic regimens was assessed based on the antibiotic susceptibility profiles of isolated bacteria. In total, 358 cases of microbiologically proven HVO were identified. The main causative pathogens identified were methicillin-susceptible *Staphylococcus aureus* (33.5%), followed by methicillin-resistant *S. aureus* (MRSA) (24.9%), aerobic gram-negative bacteria (21.8%), and *Streptococcus* species (11.7%). Extended spectrum β-lactamase (ESBL)-producing *Enterobacteriaceae* and anaerobes accounted for only 1.7% and 1.4%, respectively, of the causative pathogens. Based on the susceptibility results of isolated organisms, levofloxacin plus rifampicin was appropriate in 73.5%, levofloxacin plus clindamycin in 71.2%, and amoxicillin-clavulanate plus ciprofloxacin in 64.5% of cases. These oral combinations were more appropriate for treating community-acquired HVO (85.8%, 84.0%, and 80.4%, respectively) than healthcare-associated HVO (52.6%, 49.6%, and 37.6%, respectively). Vancomycin combined with ciprofloxacin, ceftriaxone, ceftazidime, or cefepime was similarly appropriate (susceptibility rates of 93.0%, 94.1%, 95.8%, and 95.8%, respectively). In conclusion, in a setting with a high prevalence of MRSA HVO, oral antibiotic combinations may be suboptimal for treatment of culture-negative HVO and should be used only in patients with community-acquired HVO. Vancomycin combined with fluoroquinolone or a broad-spectrum cephalosporin was appropriate in most cases of HVO in this study.

---

The incidence of vertebral osteomyelitis has increased over recent years, likely due to longer life expectancies, higher prevalence of chronic disease, and more effective diagnostic techniques (1–4). Furthermore, healthcare-associated infections, such as catheter-related and device-related bloodstream infections, also increase the risk of vertebral osteomyelitis (5, 6). Identification of microorganisms is crucial to guide appropriate antibiotic therapy for vertebral osteomyelitis, but causative microorganisms are frequently not identified. Some authors recommended a second percutaneous needle biopsy in patients with negative blood culture and initial biopsy results, but a microbiological diagnosis was still not achieved in almost one-quarter of cases (7, 8). The incidence of vertebral osteomyelitis without microbiological diagnosis has increased in recent years (1, 3). However, the optimal empiric antibiotic regimens for culture-negative vertebral osteomyelitis have yet to be determined.

The selection of an empiric antimicrobial regimen for vertebral osteomyelitis with no microbiological diagnosis should be based on consideration of the most likely causative pathogens and knowledge of local susceptibility patterns. The aim of this study was to evaluate the appropriateness of empiric antibiotic regimens suggested for treatment of culture-negative hematogenous vertebral osteomyelitis (HVO) using the susceptibility data of pathogens isolated from cases of culture-proven HVO. We hypothesized that patients with healthcare-associated HVO would more frequently be infected with antibiotic-resistant organisms than those with community-acquired HVO. Therefore, the appropriateness of empiric antibiotic regimens was also assessed according to the risk of healthcare-associated infection.

## METHODS

### Study design and setting

This observational cohort study was undertaken at five tertiary-care hospitals in the Republic of Korea, and included all adult patients diagnosed with HVO from January 2005 to December 2012. The study was conducted using the format recommended by the Strengthening the Reporting of Observational Studies in Epidemiology (STROBE) guidelines (9). This study was approved by the Institutional Review Board of Gyeongsang National University Changwon Hospital (GNUCH 2018-05-010).

### Inclusion and exclusion criteria

Adult patients (≥ 16 years of age) who presented with microbiologically proven HVO were included in this study. The diagnosis of HVO was established using previously published criteria (10, 11), which included compatible clinical features, radiological evidence of vertebral osteomyelitis, and microbiologic demonstration of bacterial pathogens, either from the site of infection itself *(e.g*., abscess, intervertebral disc, or vertebral bone) or in the blood. Possible skin contaminants, such as coagulase-negative staphylococci (CoNS) and *Propionibacterium acnes*, were considered as true pathogens if they were isolated from ≥ 2 surgical, percutaneous biopsy, or blood cultures. Cases were excluded if there was a nonhematogenous source of vertebral infection, which included (1) penetrating trauma, (2) previously placed hardware, (3) laminectomy within 1 year prior to the vertebral osteomyelitis diagnosis, or (4) the presence of a stage 3-4 decubitus ulcer at the time of diagnosis (10, 11). Cases of tuberculous, brucellar, and fungal vertebral osteomyelitis were excluded.

### Data collection

Medical records were reviewed retrospectively for demographic information, underlying illness/conditions, presumed source of infection, diagnostic work-up, clinical presentation, and laboratory and radiological data. The present study builds on our previous work on the optimal duration of antibiotic therapy in patients with HVO (10).

### Microbiological analysis

The species and susceptibility profiles of all isolates were confirmed using either the Vitek2 (bioMérieux, Marcy-l’Etoile, France) or Microscan (Dade Behring Inc., Deerfield, IL) automated systems. The susceptibility to 12 antibiotics, alone or in combination, was assessed (amoxicillin-clavulanate, ciprofloxacin, levofloxacin, clindamycin, rifampin, trimethoprim-sulfamethoxazole, fusidic acid, cefazolin, ceftriaxone, ceftazidime, cefepime, and vancomycin). Isolated strains were categorized by the Clinical Laboratory Standard Institute (CLSI) guidelines as susceptible, intermediate, or resistant to antimicrobial agents (12). Given the risk of treatment failure for intermediate strains, strains classified “intermediate” were reclassified as resistant. If the susceptibility results for surgery, percutaneous biopsy, and blood cultures were different, overall susceptibility was classified resistant.

The appropriateness of eight antibiotics, alone or in combination, for empiric treatment of HVO were analyzed: three fluoroquinolone-based oral combinations (levofloxacin plus rifampin, levofloxacin plus clindamycin, ciprofloxacin plus amoxicillin-clavulanate), four vancomycin-based intravenous combinations (vancomycin plus ciprofloxacin, ceftriaxone, ceftazidime, and cefepime), and cefazolin monotherapy. For *Staphylococcus* species, overall susceptibility to fluoroquinolone-based combinations was classified as susceptible if the organisms were susceptible to both antimicrobial agents (8, 13), except for clindamycin. For all other species, strains were considered susceptible if they showed susceptibility to at least one of the antimicrobial agent combinations.

### Definitions

HVO was classified as community-acquired or healthcare-associated HVO according to published criteria (5). Healthcare-associated HVO was defined as (i) onset of symptoms after 1 month of hospitalization with no evidence of vertebral osteomyelitis at admission, (ii) hospital admission within 6 months before symptom onset, or (iii) ambulatory diagnostic or therapeutic manipulations within 6 months before symptom onset (long-term central venous catheter use, arteriovenous fistula for hemodialysis, invasive intravascular techniques, urological, gynecological or digestive procedures, and cutaneous manipulations). Cases that did not meet any of the above criteria were classified as community-acquired HVO (5).

### Statistical analyses

Categorical variables are expressed as numbers and percentages and were compared by *χ*^2^ or Fisher’s exact test. Continuous variables are expressed as medians and interquartile ranges and were compared using Mann-Whitney *U* test. All statistical tests were two-tailed, and *P* ≤ 0.05 was considered to indicate statistical significance. All statistical analyses were performed using SPSS for Windows software (ver. 18.0; SPSS, Inc., Chicago, IL).

## RESULTS

In total, 370 patients with microbiologically proven HVO were identified during the study period. Of the 370 cases, 12 were excluded due to incomplete medical records and 358 were included in the final analysis.

### Patients’ characteristics

The demographic and baseline characteristics of the patients are displayed in Table 1. The median age of the cohort was 65 years, and 186 (52.0%) were males. The most common underlying condition was diabetes (29.3%), followed by liver cirrhosis (9.2%) and malignancy (8.7%). According to the predefined criteria, 225 (62.8%) cases were community-acquired HVO and 133 (37.2%) were healthcare-associated HVO.

**Table 1.** Demographic and baseline characteristics of 358 patients with hematogenous vertebral osteomyelitis

| Variable | All patients<br>(*n* = 358) |
| --- | --- |
| Age, median years (IQR) | 65 (58–72) |
| Male sex | 186 (52.0) |
| Underlying illness/conditions |  |
| Diabetes mellitus | 105 (29.3) |
| Liver cirrhosis | 33 (9.2) |
| Malignancy | 31 (8.7) |
| Immunosuppression | 20 (5.6) |
| End-stage renal disease | 15 (4.2) |
| Rheumatic disease | 12 (3.4) |
| Healthcare-associated HVO | 133 (37.2) |
| Presumed source of infection |  |
| Urinary tract | 35 (9.7) |
| Skin and subcutaneous tissues | 30 (8.4) |
| Intra-abdomen | 26 (7.3) |
| Infected vascular access | 25 (7.0) |
| Endocarditis | 19 (5.3) |
| Unknown | 223 (62.3) |
| Clinical data |  |
| Time to diagnosis, median days (IQR) | 22 (8–40) |
| Back pain | 319 (89.1) |
| Body temperature > 38°C | 190 (53.1) |
| Neurologic deficit at diagnosis | 61 (17.0) |
| Concurrent metastatic infection | 46 (12.8) |
| Laboratory data |  |
| White blood cell count, $\times 10^9/L$, median (IQR) | 114 (79–162) |
| C-reactive protein, mg/dL, median (IQR) | 13 (6–22) |
| Erythrocyte sedimentation rate, mm/h, median (IQR)<sup>a</sup> | 76 (55–100) |
| Positive blood cultures | 265/339 (78.2) |
| Radiologic data |  |
| Involvement of > 2 vertebral bodies | 125 (34.9) |
| Involvement of cervical spine | 31 (8.7) |
| Involvement of thoracic spine | 85 (23.7) |
| Involvement of lumbosacral spine | 283 (79.1) |
| Epidural involvement<sup>b</sup> | 194 (54.2) |
| Paravertebral involvement<sup>b</sup> | 192 (53.6) |
| Psoas involvement<sup>b</sup> | 126 (35.2) |

Data are no. (%) of patients, unless otherwise indicated.
Abbreviations: HVO, hematogenous vertebral osteomyelitis; IQR, interquartile range
<sup>a</sup> Measured in 287 patients.
<sup>b</sup> Either abscess or phlegmon.

### Microbiological findings

Of the 358 cases with microbiologically proven HVO, 93 (26.0%) were identified by diagnostic biopsy, 173 (48.3%) by blood cultures, and 92 (25.7%) by both. The most frequently isolated organisms were methicillin-susceptible *Staphylococcus aureus* (MSSA) (33.5%), follow by methicillin-resistant *S. aureus* (MRSA [24.9%]), aerobic gram-negative bacteria (21.8%), and *Streptococcus* species (11.7%; Table 2). Of 78 aerobic gram-negative bacteria, *Enterobacteriaceae* was responsible for 69 (88.5%) cases and other gram-negative bacteria for 9 (11.5%) cases. Of 42 *Streptococcus* species, viridans group streptococci were the most frequently isolated organisms (50.0%), followed by *Streptococcus agalactiae* (35.7%), and *S. pneumoniae* (9.5%). CoNS, *Enterococcus* species, and anaerobes accounted for 2.8%, 2.8%, and 1.4%, respectively.

**Table 2.** Antibiotic susceptibility testing results for 358 isolated microorganisms

|  | Percentage of isolates susceptible to |  |  |  |  |  |  |  |  |  |  |  |
| --- | --- | --- | --- | --- | --- | --- | --- | --- | --- | --- | --- | --- |
|  | AMX-CL | CIP | LEVO | CLM | RIF | TMP-SMX | FA<sup>a</sup> | CFZ | CTR | CAZ | FEP | VAN |
| Methicillin-susceptible *S. aureus* (*n* = 120) | 100 | 94.2 | 95.8 | 88.3 | 97.5 | 100 | 74.1 | 100 | NA | NA | NA | 100 |
| Methicillin-resistant *S. aureus* (*n* = 89) | 0 | 53.9 | 56.2 | 41.6 | 77.5 | 92.1 | 81.2 | 0 | NA | NA | NA | 100 |
| Gram-negative bacteria (*n* = 78) | 69.2 | 76.9 | 76.9 | NA | NA | 76.9 | NA | 80.8 | 82.1 | 89.7 | 89.7 | NA |
| *Streptococcus* species (*n* = 42) | 95.2 | NA | 90.5 | 76.2 | NA | NA | NA | 100<sup>b</sup> | 95.2 | NA | NA | 100 |
| Coagulase-negative staphylococci (*n* = 10) | 50.0 | 70.0 | 70.0 | 70.0 | 80.0 | 80.0 | NA | 50.0 | NA | NA | NA | 100 |
| *Enterococcus* species (*n* = 10) | 90.0 | NA | NA | NA | NA | NA | NA | 0 | NA | NA | NA | 100 |
| Other (*n* = 9)<sup>c</sup> | 55.6 | 11.1 | 11.1 | NA | NA | NA | NA | 0 | 11.1 | 11.1 | 11.1 | NA |

Abbreviations: AMX-CL, amoxicillin-clavulanate; CFP, cefepime; CAZ, ceftazidime; CFZ, cefazolin; CIP, ciprofloxacin; CLM, clindamycin;
CTR, ceftriaxone; FA, fusidic acid; LEVO, levofloxacin; MRSA, methicillin-resistant *S. aureus*; MSSA, methicillin-susceptible *S. aureus*;
RIF, rifampin; TMP-SMX; trimethoprim-sulfamethoxazole; VANC, vancomycin.
<sup>a</sup> Susceptibility data for fusidic acid against methicillin-susceptible *S. aureus* and methicillin-resistant *S. aureus* isolates were available in 112
and 85 cases, respectively.
<sup>b</sup> Excluding 25 streptococcal isolates without susceptibility data (21 viridans group streptococci and 4 *S. pneumoniae*); the remaining 17
*Streptococcus* isolates were susceptible to cefazolin.
<sup>c</sup> Included *Bacteroides fragilis* ( $n = 2$ ), *Bacteroides ureolyticus* ( $n = 1$ ), *Prevotella melaninogenica* ( $n = 1$ ), *Prevotella oralis* ( $n = 1$ ),
*Micrococcus* species ( $n = 1$ ), *Enterococcus faecium*/viridans group streptococci ( $n = 1$ ), *Klebsiella pneumoniae*/*Enterobacter aerogenes* ( $n =$
1), and methicillin-susceptible *S. aureus*/methicillin-resistant *Staphylococcus schleiferi* ( $n = 1$ ).

There were some differences in the proportions of pathogens between community-acquired HVO and healthcare-associated HVO. MRSA was more frequent in healthcare-associated HVO than in community-acquired HVO (43.6% vs. 13.8%; *P* < 0.001), but MSSA was more frequent in community-acquired HVO than in healthcare-associated HVO (44.0% vs. 13.8%; *P* < 0.001). *Streptococcus* species were more frequent in community-acquired cases than in healthcare-associated cases (16.0% vs. 4.5%; *P* = 0.001). The rate of blood culture-proven cases among all culture-proven cases was 81.7% in *S. aureus* HVO, 71.8% in gram-negative bacterial HVO, and 61.9% in streptococcal HVO.

### Antimicrobial susceptibility profiles of isolated organisms

The antimicrobial susceptibility testing results of isolated microorganisms are shown in Table 2. About 95% of MSSA isolates were susceptible to fluoroquinolones, compared to only half of the MRSA isolates. Of 69 *Enterobacteriaceae* isolates, 18 (26.1%) were resistant to ciprofloxacin, and 6 (8.7%) were extended-spectrum β-lactamase (ESBL) producers. All of the remaining nine gram-negative bacteria, including five *Pseudomonas aeruginosa* isolates, were susceptible to ciprofloxacin. Of 42 *Streptococcus* species isolates, 40 (95.2%) were susceptible to amoxicillin-clavulanate and 38 (90.5%) were susceptible to levofloxacin. Of the 10 CoNS isolates, 5 (50%) were methicillin-resistant.

### Relevance of empirical antibiotic regimens for culture-negative HVO

The relevance of empiric antibiotic regimens for culture-negative HVO is shown in Table 3. Based on the susceptibility results of isolated organisms, levofloxacin plus rifampicin was appropriate in 73.5% of cases, levofloxacin plus clindamycin in 71.2% of cases, and amoxicillin-clavulanate plus ciprofloxacin in 64.5% of cases. These oral combinations were more appropriate for community-acquired HVO (85.8%, 84.0%, and 80.4%, respectively) than healthcare-associated HVO (52.6%, 49.6%, and 37.6%, respectively). After excluding 25 cases of α-hemolytic streptococcal HVO without cefazolin susceptibility test results, cefazolin was appropriate in 61.6% of HVO cases. Cefazolin was appropriate in 78.3% of community-acquired cases and 35.4% of healthcare-associated cases. Vancomycin combined with ciprofloxacin, ceftriaxone, ceftazidime, or cefepime was similarly appropriate (susceptibility rates 93.0%, 94.1%, 95.8%, and 95.8%, respectively). All antibiotic regimens gave similar results, irrespective blood culture positivity.

**Table 3.** Relevance of empiric antibiotic therapy based on the susceptibility results of organisms isolated from cases of culture-proven hematogenous vertebral osteomyelitis

| Organism | LEVO +<br>RIF | LEVO +<br>CLM | AMX-CL<br>+ CIP | CFZ<sup>a</sup> | VAN +<br>CIP | VAN +<br>CTR | VAN +<br>CAZ | VAN +<br>FEP |
| --- | --- | --- | --- | --- | --- | --- | --- | --- |
| All cases (*n* = 358) | 73.5 | 71.2 | 66.5 | 61.6 | 93.0 | 94.1 | 95.8 | 95.8 |
| Microorganisms |  |  |  |  |  |  |  |  |
| Methicillin-susceptible *S. aureus* (*n* = 120) | 93.3 | 88.3 | 100 | 100 | 100 | 100 | 100 | 100 |
| Methicillin-resistant *S. aureus* (*n* = 89) | 50.6 | 41.6 | 0 | 0 | 100 | 100 | 100 | 100 |
| Gram-negative bacteria (*n* = 78) | 76.9 | 76.9 | 71.8 | 80.8 | 76.9 | 82.1 | 89.7 | 89.7 |
| *Streptococcus* species (*n* = 42) | 90.5 | 97.6 | 100 | 100<sup>a</sup> | 100 | 100 | 100 | 100 |
| Coagulase-negative staphylococci (*n* = 10) | 70.0 | 70.0 | 50.0 | 50 | 100 | 100 | 100 | 100 |
| *Enterococcus* species (*n* = 10) | 0 | 0 | 90.0 | 0 | 100 | 100 | 100 | 100 |
| Other (*n* = 9) | 11.1 | 44.4 | 66.7 | 0 | 22.2 | 22.2 | 22.2 | 22.2 |
| Onset of infection |  |  |  |  |  |  |  |  |
| Community-acquired (*n* = 225) | 85.8 | 84.0 | 80.4 | 78.3 | 94.2 | 96.9 | 97.3 | 97.3 |
| Healthcare-associated (*n* = 133) | 52.6 | 49.6 | 37.6 | 35.4 | 91.0 | 89.5 | 93.2 | 93.2 |
| Blood culture |  |  |  |  |  |  |  |  |
| Non-bacteremic HVO (*n* = 93)<sup>b</sup> | 77.4 | 79.6 | 66.7 | 61.4 | 91.7 | 86.0 | 92.5 | 92.5 |
| Bacteremic HVO (*n* = 265) | 72.1 | 68.3 | 63.8 | 61.6 | 92.8 | 97.0 | 97.0 | 97.0 |

Data are no. (%) of isolates, unless otherwise indicated.
Abbreviations: AMX-CL, amoxicillin-clavulanate; FEP, cefepime; CAZ, ceftazidime; CIP,
ciprofloxacin; CFZ, cefazolin; CLM, clindamycin; CTR, ceftriaxone; HVO, hematogenous
vertebral osteomyelitis; LEVO, levofloxacin; RIF, rifampin; VAN, vancomycin.
<sup>a</sup> Excluding 25 streptococcal isolates without susceptibility data (21 viridans group
streptococci and 4 *S. pneumoniae*); the remaining 17 *Streptococcus* isolates were susceptible
to cefazolin.
<sup>b</sup> Included patients with negative blood culture results ( $n = 74$ ) and those from whom blood cultures were not obtained ( $n = 19$ ).

## DISCUSSION

We evaluated the susceptibility profiles of microorganisms causing HVO and assessed the appropriateness of several empiric antibiotic regimens for culture-negative HVO. Our data showed that oral antibiotic combinations may be suboptimal for empiric treatment of culture-negative HVO, and should be used after definitively ruling out healthcare-associated HVO. Vancomycin combined with fluoroquinolones or broad-spectrum cephalosporins were appropriate for most cases of community-acquired and healthcare-associated HVO.

In this study, the most frequently isolated organism was *S. aureus*, with 43% being MRSA. In recent years, the prevalence of MRSA among *S. aureus* vertebral osteomyelitis has been reported to be 40-57% (11, 14–16). The resistance rate of *Enterobacteriaceae* isolates to ciprofloxacin was 26% in this study, compared to 31% and 38% in two recent studies conducted in South Korea and France, respectively (17, 18). Vertebral osteomyelitis caused by antibiotic-resistant organisms may have a higher risk of treatment failure (16, 19, 20). Indeed, patients with MRSA vertebral osteomyelitis reportedly have a 4-5-fold higher risk of recurrence than those with MSSA vertebral osteomyelitis (16, 19). In a study of patients with gram-negative bacterial HVO, half of those infected with fluoroquinolone-resistant strains experienced recurrence after 4-8 weeks of antibiotic treatment (20). Thus, the increasing incidence of antibiotic resistance in causative pathogens of vertebral osteomyelitis should be considered when selecting an empiric antibiotic regimen for culture-negative HVO.

Fluoroquinolone-based regimens have high oral bioavailability and good bone penetration, allow an early switch to oral therapy, and appear to be the best option for empiric treatment of culture-negative vertebral osteomyelitis (8, 21). Two previous studies showed that empiric treatment with levofloxacin plus rifampin had overall response rates of 78% (3) and 84% (22). In another study, empiric treatment with ciprofloxacin plus amoxicillin-clavulanate had an overall response rate of 82% (23). All three studies were conducted in an epidemiologicsetting featuring a low prevalence of MRSA (3, 22, 23), and the appropriateness of these regimens in high-MRSA-prevalence settings was not evaluated. In the current study, based on the susceptibility profiles of isolated bacteria, these combination regimens were appropriate in 74% and 67% of cases, respectively. We also found that levofloxacin plus clindamycin exhibited activity against 71% of causative organisms of HVO. Fluoroquinolone plus clindamycin is suggested for the treatment of culture-negative vertebral osteomyelitis by British guidelines (4), but clinical data on this regimen in this setting are limited. The marked inappropriateness of fluoroquinolone-based regimens in the current study may be due to the high prevalence of MRSA (25%). Almost half of our MRSA isolates were resistant to fluoroquinolones, and the resistance rates to their companion drugs, such as rifampin and clindamycin, were high (22% and 58%, respectively). Recently, Desoutter *et al*. reported the susceptibility patterns of microorganisms isolated from 101 cases of non-bacteremic vertebral osteomyelitis by percutaneous needle biopsy (17). According to the antibiotic susceptibility profiles, fluoroquinolone-based regimens were appropriate in 58-77% of cases (17). In contrast to our study, MRSA was responsible for only one case of vertebral osteomyelitis, but CoNS was the most frequent isolate, with 20% of isolates being resistant to fluoroquinolones (17). Also, the almost 40% rate of fluoroquinolone resistance in *Enterobacteriaceae* isolates contributed to the relatively low suitability of fluoroquinolone-based regimens (17). To minimize the risk of microbiological failure, we reassessed the suitability of empiric regimens in community-acquired and healthcare-associated cases. These analyses indicated that fluoroquinolone-based regimens were appropriate in up to 86% of community-acquired HVO cases, but in only 38-53% of healthcare-associated HVO cases. Thus, based on current data, as well as previous clinical experience (3, 22, 23), fluoroquinolone-based regimens may be useful for empiric treatment of community-acquired HVO, but they should not be used for healthcare-associated HVO.

In this study, vancomycin plus ciprofloxacin or broad-spectrum cephalosporins were appropriate in most cases of community-acquired and healthcare-associated HVO. Although ESBL-producing *Enterobacteriaceae* accounted for 9% of the *Enterobacteriaceae* isolates, it accounted for only 2% of all causative organisms. The prevalence of anaerobes was 1% in this work, and up to 4% in two recent studies (3, 21). Thus, antibiotic coverage of such rare organisms (with carbapenems or metronidazole) may be not required initially, and should be limited to patients who show no clinical improvement after empiric treatment and negative results of repeated biopsies. As a companion drug to vancomycin, fluoroquinolones and broad-spectrum cephalosporins were similarly effective in healthcare-associated HVO. Taking into consideration the bone penetration of antibiotics (24), we prefer fluoroquinolones over broad-spectrum cephalosporins as companion drugs to vancomycin.

For culture-negative vertebral osteomyelitis, some authors suggest a first-generation cephalosporin for community-acquired cases and vancomycin-containing regimens for post-surgical cases, and have reported favorable outcomes with this approach (25, 26). A first-generation cephalosporin is an acceptable alternative in patients unable to tolerate anti-staphylococcal penicillin (27), but clinical experience with this agent for streptococcal and gram-negative bacterial osteomyelitis is limited. In addition, first-generation cephalosporins show varying activity against α-hemolytic streptococci, unlike P-hemolytic streptococci (28), and susceptibility testing of this agent against α-hemolytic streptococci is not recommended (12). Even after excluding 25 cases of α-hemolytic streptococci without susceptibility data (viridans group streptococci [n = 21] and *S. pneumoniae* [n = 4]), cefazolin was appropriate in only 62% of cases. Given its poor penetration of bone tissue, oral cephalosporin is not recommended for the treatment of vertebral osteomyelitis by the Infectious Disease Society of America (IDSA) (21) and French (13) guidelines. More data on the efficacy of first-generation cephalosporins are required before they can be widely used for culture-negative vertebral osteomyelitis.

Our study had several limitations. First, some cases lacked antibiotic susceptibility test results and so exclusion of these cases may have introduced bias. Second, we assessed the appropriateness of empiric antibiotic regimens for culture-negative HVO based on the data of organisms isolated from microbiologically proven cases. However, culture-negative vertebral osteomyelitis may be more closely related to cases caused by low-virulence pathogens, such streptococci and CoNS (3, 29), and the prevalence of these organisms may have been underestimated in our analysis. In this study, *Streptococcus* species were less likely to be detected in blood cultures than *S. aureus* (62% vs. 82%). Finally, this study included only patients with HVO, and so our findings should not be extrapolated to cases of post-surgical vertebral osteomyelitis, in which CoNS and antibiotic-resistant organisms may be more common.

In summary, in a setting in which MRSA is a frequent causative organism of HVO, cautious selection of empiric antibiotics is required. Oral empiric antibiotic combinations may be suboptimal for culture-negative HVO, and should be used only in community-acquired cases. Vancomycin combined with fluoroquinolone or a broad-spectrum cephalosporin is appropriate in most cases of HVO. The clinical efficacy of empiric antibiotic regimens for culture-negative HVO reported here should be verified in further studies.

## References

1. Kehrer M, Pedersen C, Jensen TG, and Lassen AT. 2014. Increasing incidence of pyogenic spondylodiscitis: a 14-year population-based study. J. Infect. 68: 313–320.

2. Akiyama T, Chikuda H, Yasunaga H, Horiguchi H, Fushimi K, and Saita K. 2013. Incidence and risk factors for mortality of vertebral osteomyelitis: a retrospective analysis using the Japanese diagnosis procedure combination database. BMJ Open 3.

3. Lora-Tamayo J, Euba G, Narvaez JA, Murillo O, Verdaguer R, Sobrino B, Narvaez J, Nolla JM, and Ariza J. 2011. Changing trends in the epidemiology of pyogenic vertebral osteomyelitis: the impact of cases with no microbiologic diagnosis. Semin. Arthritis Rheum. 41: 247–255.

4. Cottle L, and Riordan T. 2008. Infectious spondylodiscitis. J. Infect. 56: 401–412.

5. Pigrau C, Rodriguez-Pardo D, Fernandez-Hidalgo N, Moreto L, Pellise F, Larrosa MN, Puig M, and Almirante B. 2015. Health care associated hematogenous pyogenic vertebral osteomyelitis: a severe and potentially preventable infectious disease. Medicine (Baltimore) 94:e365.

6. Renz N, Haupenthal J, Schuetz MA, and Trampuz A. 2017. Hematogenous vertebral osteomyelitis associated with intravascular device-associated infections - A retrospective cohort study. Diagn Microbiol Infect Dis 88: 75–81.

7. Gras G, Buzele R, Parienti JJ, Debiais F, Dinh A, Dupon M, Roblot F, Mulleman D, Marcelli C, Michon J, and Bernard L. 2014. Microbiological diagnosis of vertebral osteomyelitis: relevance of second percutaneous biopsy following initial negative biopsy and limited yield of post-biopsy blood cultures. Eur J Clin Microbiol Infect Dis 33: 371–375.

8. Berbari EF, Kanj SS, Kowalski TJ, Darouiche RO, Widmer AF, Schmitt SK, Hendershot EF, Holtom PD, Huddleston PM, 3rd, Petermann GW, and Osmon DR. 2015. 2015 Infectious Diseases Society of America (IDSA) Clinical Practice Guidelines for the Diagnosis and Treatment of Native Vertebral Osteomyelitis in Adults. Clin Infect Dis 61:e26–46.

9. von Elm E, Altman DG, Egger M, Pocock SJ, Gotzsche PC, Vandenbroucke JP, and Initiative S. 2007. The Strengthening the Reporting of Observational Studies in Epidemiology (STROBE) statement: guidelines for reporting observational studies. Lancet 370: 1453–1457.

10. Park KH, Cho OH, Lee JH, Park JS, Ryu KN, Park SY, Lee YM, Chong YP, Kim SH, Lee SO, Choi SH, Bae IG, Kim YS, Woo JH, and Lee MS. 2016. Optimal Duration of Antibiotic Therapy in Patients With Hematogenous Vertebral Osteomyelitis at Low Risk and High Risk of Recurrence. Clin Infect Dis 62: 1262–1269.

11. Livorsi DJ, Daver NG, Atmar RL, Shelburne SA, White AC, Jr., and Musher DM. 2008. Outcomes of treatment for hematogenous *Staphylococcus aureus* vertebral osteomyelitis in the MRSA ERA. J. Infect. 57: 128–131.

12. Clinical and Laboratory Standards Institute. 2011. Performance Standards for Antimicrobial Susceptibility Testing, 21th Infomational supplment. CLSI document M100-S21. Clinical and Laboratory Standards Institue, Wayne, PA, USA.

13. Société de Pathologies Infectieuses de Langue Française (SPILF). 2007. Primary infectious spondylitis, and following intradiscal procedure, without prothesis. Recommendations. Med. Mal. Infect. 37: 573–583.

14. Bhavan KP, Marschall J, Olsen MA, Fraser VJ, Wright NM, and Warren DK. 2010. The epidemiology of hematogenous vertebral osteomyelitis: a cohort study in a tertiary care hospital. BMC Infect Dis 10: 158.

15. Weissman S, Parker RD, Siddiqui W, Dykema S, and Horvath J. 2014. Vertebral osteomyelitis: retrospective review of 11 years of experience. Scand. J. Infect. Dis. 46: 193–199.

16. Inoue S, Moriyama T, Horinouchi Y, Tachibana T, Okada F, Maruo K, and Yoshiya S. 2013. Comparison of clinical features and outcomes of *staphylococcus aureus* vertebral osteomyelitis caused by methicillin-resistant and methicillin-sensitive strains. SpringerPlus 2: 283.

17. Desoutter S, Cottier JP, Ghout I, Issartel B, Dinh A, Martin A, Carlier R, Bernard L, and Duration of Treatment for Spondylodiscitis Study G. 2015. Susceptibility Pattern of Microorganisms Isolated by Percutaneous Needle Biopsy in Nonbacteremic Pyogenic Vertebral Osteomyelitis. Antimicrob Agents Chemother 59: 7700–7706.

18. Kang SJ, Jang HC, Jung SI, Choe PG, Park WB, Kim CJ, Song KH, Kim ES, Kim HB, Oh MD, Kim NJ, and Park KH. 2015. Clinical characteristics and risk factors of pyogenic spondylitis caused by gram-negative bacteria. PLoS One 10:e0127126.

19. Park KH, Chong YP, Kim SH, Lee SO, Choi SH, Lee MS, Jeong JY, Woo JH, and Kim YS. 2013. Clinical characteristics and therapeutic outcomes of hematogenous vertebral osteomyelitis caused by methicillin-resistant *Staphylococcus aureus*. J. Infect. 67: 556–564.

20. Park KH, Cho OH, Jung M, Suk KS, Lee JH, Park JS, Ryu KN, Kim SH, Lee SO, Choi SH, Bae IG, Kim YS, Woo JH, and Lee MS. 2014. Clinical characteristics and outcomes of hematogenous vertebral osteomyelitis caused by gram-negative bacteria. J. Infect. 69: 42–50.

21. Bernard L, Dinh A, Ghout I, Simo D, Zeller V, Issartel B, Le Moing V, Belmatoug N, Lesprit P, Bru JP, Therby A, Bouhour D, Denes E, Debard A, Chirouze C, Fevre K, Dupon M, Aegerter P, Mulleman D, and Duration of Treatment for Spondylodiscitis study g. 2015. Antibiotic treatment for 6 weeks versus 12 weeks in patients with pyogenic vertebral osteomyelitis: an open-label, non-inferiority, randomised, controlled trial. Lancet 385: 875–882.

22. Viale P, Furlanut M, Scudeller L, Pavan F, Negri C, Crapis M, Zamparini E, Zuiani C, Cristini F, and Pea F. 2009. Treatment of pyogenic (non-tuberculous) spondylodiscitis with tailored high-dose levofloxacin plus rifampicin. Int. J. Antimicrob. Agents 33: 379–382.

23. Luzzati R, Giacomazzi D, Danzi MC, Tacconi L, Concia E, and Vento S. 2009. Diagnosis, management and outcome of clinically-suspected spinal infection. J. Infect. 58: 259–265.

24. Landersdorfer CB, Bulitta JB, Kinzig M, Holzgrabe U, and Sorgel F. 2009. Penetration of antibacterials into bone: pharmacokinetic, pharmacodynamic and bioanalytical considerations. Clin. Pharmacokinet. 48: 89–124.

25. Yoon SH, Chung SK, Kim KJ, Kim HJ, Jin YJ, and Kim HB. 2010. Pyogenic vertebral osteomyelitis: identification of microorganism and laboratory markers used to predict clinical outcome. Eur. Spine J. 19: 575–582.

26. Kim J, Kim YS, Peck KR, Kim ES, Cho SY, Ha YE, Kang CI, Chung DR, and Song JH. 2014. Outcome of culture-negative pyogenic vertebral osteomyelitis: comparison with microbiologically confirmed pyogenic vertebral osteomyelitis. Semin. Arthritis Rheum. 44: 246–252.

27. Lew DP, and Waldvogel FA. 2004. Osteomyelitis. Lancet 364: 369–379.

28. Fraimow HS. 2009. Systemic antimicrobial therapy in osteomyelitis. Semin. Plast. Surg. 23: 90–99.

29. Murillo O, Roset A, Sobrino B, Lora-Tamayo J, Verdaguer R, Jimenez-Mejias E, Nolla JM, Colmenero Jde D, and Ariza J. 2014. Streptococcal vertebral osteomyelitis: multiple faces of the same disease. Clin Microbiol Infect 20:O33–38.

